# Abnormal nitration and S-sulfhydration of Sp1-CSE-H_2_S pathway contribute to the progress of hyperhomocysteinemia: a vicious circle

**DOI:** 10.1101/843854

**Authors:** Chenghua Luo, Dengyu Ji, Yan Li, Yan Cao, Shangyue Zhang, Wenjing Yan, Ke Xue, Jiayin Chai, Ye Wu, Huirong Liu, Wen Wang

**Affiliations:** Department of Physiology and Pathophysiology, School of Basic Medical Sciences, Capital Medical University, Beijing 100069, China; Beijing Key Laboratory for Metabolic Disorder-Related Cardiovascular Diseases, Beijing 100069, China; Department of Pain Management, Xuanwu Hospital, Capital Medical University, Beijing, 100053, China; Center for Anesthesiology, Beijing Anzhen Hospital, Capital Medical University, Beijing, 100029, China

**Keywords:** nitration, S-sulfhydration, Specificity protein 1, Cystathionine γ-Lyase, hydrogen sulfide, hyperhomocysteinemia

## Abstract

Sp1 (Specificity protein 1)-CSE (cystathionine-γ-lyase)-H_2_S (hydrogen sulfide) pathway plays an important role in homocysteine-metabolism, whose disorder can result in hyperhomocysteinemia. The deficiency of plasma H_2_S in patients and animal models with hyperhomocysteinemia has been reported but it is unclear whether this deficiency plays a role in the progress of hyperhomocysteinemia. Furthermore, it remains unknown whether the post-translational modification of Sp1 or CSE mediated by hyperhomocysteinemia itself can in turn affect the development of hyperhomocysteinemia. By both in vivo and in vitro studies, we conducted immunoprecipitation and maleimide assays to detect the post-translational modification of Sp1-CSE-H_2_S pathway and revealed four major findings: (1) the accumulation of homocysteine augmented the nitration of CSE, thus blunted its bio-activity and caused H_2_S deficiency. (2) H_2_S deficiency lowered the S-sulfhydration of Sp1 and inhibited its transcriptional activity, resulted in lower expression of CSE. CSE deficiency decreased the H_2_S level further, which in turn lowered the S-sulfhydration level of CSE. (3) CSE was S-sulfhydrated at Cys84, Cys109, Cys172, Cys229, Cys252, Cys307 and Cys310 under physiological conditions, mutation of Cys84, Cys109, Cys229, Cys252 and Cys307 decreased its S-sulfhydration level and bio-activity. (4) H_2_S deficiency could trap hyperhomocysteinemia into a progressive vicious circle and trigger a rapid increase of homocysteine, while blocking nitration or restoring S-sulfhydration could break this circle. In conclusion, this study reveals a novel mechanism involved in the disorder of homocysteine-metabolism, which may provide a candidate therapeutic strategy for hyperhomocysteinemia.

## INTRODUCTION

As a non-essential sulfhydryl-containing amino acid derived from methionine metabolism, the elevation of serum homocysteine (Hcy) is clinically called hyperhomocysteinemia (HHcy). HHcy is an independent risk factor of cardiovascular disease [1] and associated with the onset of several neurologic disorders such as stroke [2] and Alzheimer’s disease [3]. High methionine diet, dietary deficiencies of folic acid or vitamin B, gene mutations in Hcy-metabolizing pathway [4, 5], impaired hepatic [6] and renal functions [7, 8] have been reported to be associated with HHcy. Folic acid supplementation is used to decrease plasma Hcy levels in clinic, however, it is reported that more than 40% patients with HHcy fail to reach the normal range (5-15μmol/L) after 3 months of folic acid supplementation [9]. Therefore, further prospective studies are necessary to explore new mechanism(s) involved in Hcy-metabolism and seek for novel therapeutic targets for HHcy.

HHcy induces cerebrovascular dysfunction by producing of superoxide [10]. Meanwhile, superoxide quenches nitric oxide (NO) to form peroxynitrite (ONOO^-^) which binds to tyrosine residues in proteins to produce 3-nitrotyrosine (3-NT) which consequently increases the nitration stress [11]. Our previous studies have shown that HHcy increases the nitration level of cystathionine-β-synthase (CBS), an important pyridoxal 5’-phosphate (PLP)-dependent enzyme in the trans-sulfuration pathway, inhibiting its bio-activity, which in turn accelerates the progress of HHcy [12]. Similar to CBS, CSE is another PLP-dependent enzyme that also plays an important role in the metabolism of Hcy. However, the nitration of CSE has not been reported thus far. Furthermore, it also needs to be clarified if Hcy also mediates the post-translational modification (PTM) of CSE by which affects the progress of HHcy.

H_2_S, ever regarded as a toxic gas, has recently been classified as a gasotransmitter that can modify proteins by S-sulfhydration [13–18]. In mammals, Hcy can be metabolized to generate H_2_S through the trans-sulfuration pathway [19], in which CSE is the primary enzyme that generates H_2_S in peripheral tissues [20]. It has been reported that HHcy can cause H_2_S deficiency [21–24], however, whether H_2_S deficiency can in turn affect the progress of HHcy, remains largely unknown. CSE deficiency has been reported in HHcy [25], suggesting the transcription level of *Cse* is affected. However, it is not known the impact of HHcy on its transcription factor Sp1. Further clarification is needed to understand if HHcy mediates the post-translational modification of Sp1, thus in turn affects the progress of HHcy itself.

H_2_S administration can inhibit HHcy-induced oxidative stress [26], ER stress [27], cardiac remodeling [28] and plasma lipid peroxidation [29]. Thus, administration of H_2_S could be a novel therapeutic strategy to block the injury imposed by HHcy. However, to the best of our knowledge, it has not been reported whether H_2_S can be used as a blocker during the progress of HHcy, through which avoids HHcy-induced injuries in the beginning.

Based on reported studies, we accordingly hypothesize that Hcy increases the nitration level of CSE, leading to H_2_S deficiency and subsequently affecting S-sulfhydration of Sp1-CSE-H_2_S pathway, which, in turn, influences the progress of HHcy. To test this hypothesis, we have fed mice with high methionine diet (high Met diet) and treated QSG-7701 cells with Hcy to establish an HHcy model, and examined the model by using immunoprecipitation and maleimide assays to detect the effects of Hcy on nitration and S-sulfhydration, respectively. This study aims to elucidate a novel mechanism that may be involved in the disorder of Hcy-metabolism, and therefore, provides possible therapeutic targets for HHcy.

## RESULTS

### Accumulation of Hcy augmented the nitration level of CSE

In the present study, we fed mice with high Met diet to establish HHcy model. After feeding for 8 weeks, we detected a significant increase of total serum Hcy (tHcy) in high Met diet-fed group mice compared to the wild type group (WT) (Fig 1A). Additionally, an increase of 3-NT in the liver lysate was also found (Fig 1B). With the feeding continuing to 17 weeks, the tHcy (Fig 1C) and 3-NT level (Fig 1D) increased further. In vitro, we treated QSG-7701 cells with 2mM Hcy for 24h, a significant increase of 3-NT was found and meanwhile, pretreatment with Fe (III) meso-tetra (N-methyl-4-pyridyl) porphine pentachloride (FeTMPyP), a scavenger reagent of ONOO^-^, reduced the generation of 3-NT (Fig 1E). Furthermore, immunoprecipitation detection revealed that the level of nitrated CSE in the liver of mice (fed with high Met diet) was increased significantly compared to WT (Fig 1F-G). Similarly, treating QSG-7701 cells with 2mM Hcy for 24h also led to a significant increase of nitrated CSE (Fig 1H).

**Fig 1.**
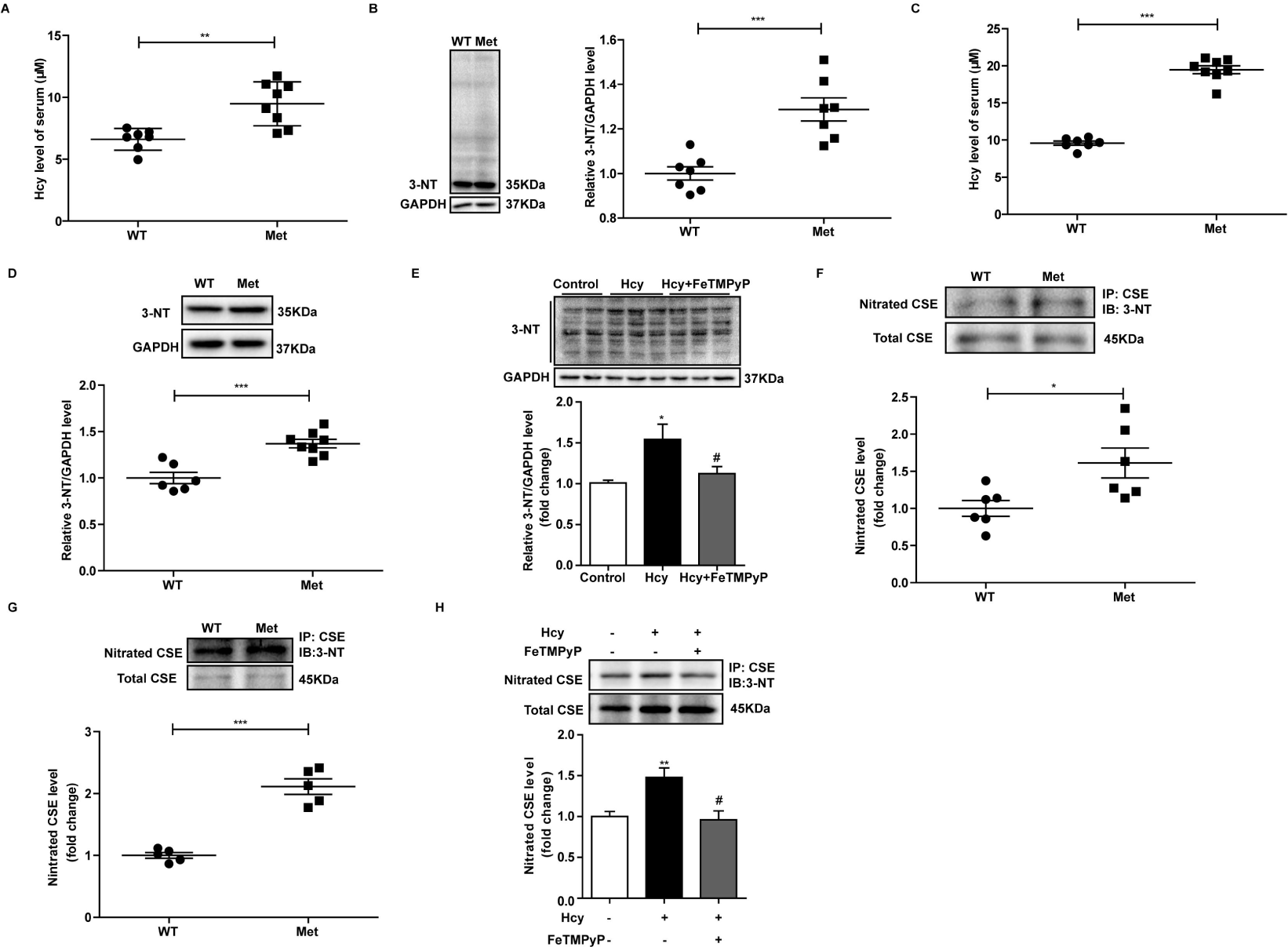
Hcy augmented the nitration level of CSE. (A) Serum Hcy level of mice fed with high Met diet for 8 weeks. (B) 3-NT level in liver lysate of mice fed with high Met diet for 8 weeks. (C) Serum Hcy level of mice fed with high Met diet for 17 weeks. (D) 3-NT level in liver lysate of mice fed with high Met diet for 17 weeks. (E) 3-NT level in QSG-7701 cells was determined by western blot, n=3. (F) The nitration level of CSE in liver lysate of mice fed with high Met diet for 8 weeks. (G) The nitration level of CSE in liver lysate of mice fed with high Met diet for 17 weeks. (H) Nitration of CSE in QSG-7701 cells was determined by immunopricipitation, n=3. Graphs show means ± SEM. [Student’s t test, (A, B, C, D, F, G), \**P*≤0.05, \*\**P*≤0.01, Met vs Control (WT). one-way ANOVA. NS (no significant), (E, H), Hcy vs Control, \**P*≤0.05, \*\**P*≤0.01, Hcy (Met) vs Control (WT). ^#^*P*≤0.05, Hcy+FeTMPyP vs Hcy.]. Cells were treated with Hcy (2mM), FeTMPyP (100μM) for 24h then harvested for detection.

These results indicated that, similar to CBS, CSE could also be nitrated under physiological conditions, and its nitration level could be augmented by the accumulation of Hcy.

### Hcy-induced nitration inhibited the bio-activity of CSE and lowered H_2_S production

Nitration generally inhibits the bio-activity of protein. To further confirm its effect on the enzymatic activity of CSE, we utilized FeTMPyP to block the augment of nitration induced by Hcy in vitro. As shown in Fig 2A, the decreased enzymatic activity of CSE induced by Hcy was rescued by FeTMPyP. Additionally, mice fed with high Met diet for 8 weeks exhibited a significant decline of CSE’s activity compared to WT (Fig 2B). These results indicated that Hcy augmented the nitration level of CSE and inhibited its bio-activity.

**Fig 2.**
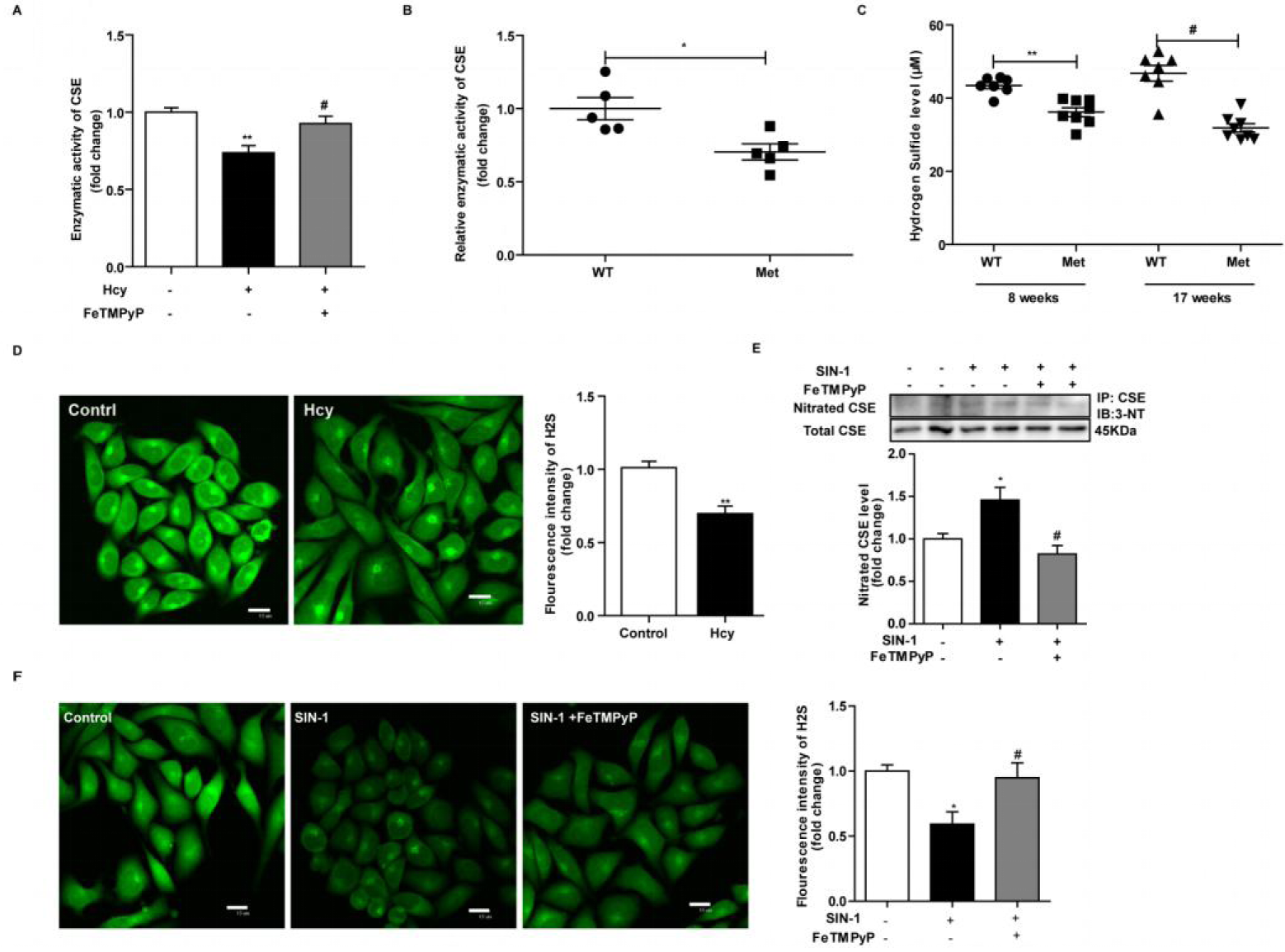
Nitration inhibited the bio-activity of CSE thus decreased the production of H_2_S. (A) Enzymatic activity of CSE in QSG-7701 cells, n=3. (B) Enzymatic activity of CSE in liver lysate of mice fed with high Met diet for 8 weeks. (C) H_2_S level in the serum of mice, determined by methylene blue assay. (D) H_2_S level in QSG-7701 cells treated with or without Hcy (2mM, 36h) detected by the H_2_S-selective sensor C7-Az, bar: 15μm, n=3. (E) QSG-7701 cells treated with SIN-1 (200μM) for 24h were harvested to detect the nitration level of CSE, n=4. (F) H_2_S was detected by C7-Az in QSG-7701 cells treated with SIN-1 (200μM) for 24h, bar: 15μm, n=3. Graphs show means ± SEM. [Student’s t test, (B, D, F), one-way ANOVA. NS (no significant), (A, C, E, F), *\*P* ≤0.05, \*\**P*≤0.01, ^#^*P*≤0.05, SIN-1 vs Control, ^#^*P*≤0.05, SIN-1+FeTMPyP vs SIN-1].

In view of it is CSE that endogenously catalyzed the production of H_2_S in mammalian liver [30–32], we then explored whether H_2_S production could be affected by Hcy. As shown in Fig 2C, mice fed with high Met diet exhibited a significant decrease of serum H_2_S level. In vitro, we used a H_2_S-selective sensor C7-Az to detect the intracellular H_2_S, and found that QSG-7701 cells treated with Hcy for 24h produced less H_2_S compared to control (Fig 2D). These results confirmed that the increase of Hcy level was correlated with the decline of H_2_S.

Additionally, to further confirm whether the increased nitration level of CSE could decrease the production of H_2_S, SIN-1, a direct ONOO^-^ donor was used in vitro. As shown in Fig 2E, after treating with SIN-1, the nitration level of CSE augmented, sequentially decreased the intracellular H_2_S level (Fig 2F), while preconditioning with FeTMPyP could block the augment of nitration and sequentially rescue the decline of H_2_S.

These results confirmed that Hcy induced augment of nitration could lower H_2_S production.

### The S-sulfhydration level and transcription activity of Sp1 was decreased in HHcy

In the liver of mice fed with high Met diet for 8 weeks, the protein level of CSE exhibited no significant change (S1A Fig). However, an obvious decline of CSE’s mRNA was detected (Fig 3A). Meanwhile, in vitro, we treated QSG-7701 cells with Hcy for 24h and detected a decline of *Cse*’s mRNA level as well (S1B Fig). With time continuing, we found a significant decrease of CSE’s protein level in mice (high Met diet for 17 weeks) (Fig 3B) and QSG-7701 cells (Hcy treated for 36h) (Fig 3C, D). These results hinted the transcription level of CSE had been affected by Hcy.

**Fig 3.**
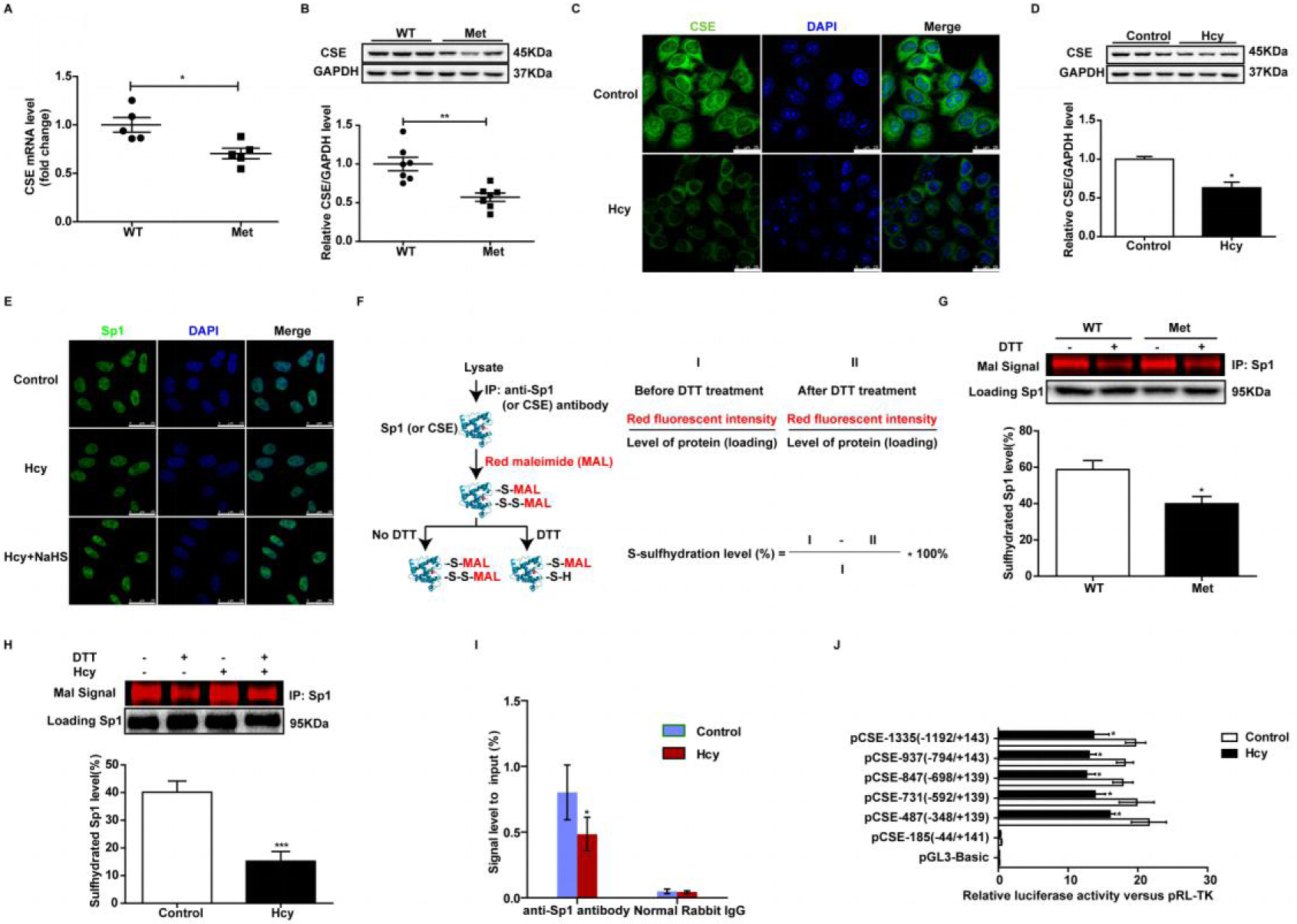
The S-sulfhydration of Sp1 was decreased in HHcy. (A) The mRNA of *Cse* in the liver of mice fed with high Met diet for 8 weeks. (B) Protein level of CSE in wild type and mice fed with high Met diet for 17 weeks. (C) Immunofluorescence detection of the sub-cellular location and expression of CSE in QSG-7701 cells, n=3. (D) The effects of Hcy (2mM, 36h) on the level of CSE in QSG-7701 cells, n=3. (E) The subcellular location of Sp1 in QSG-7701 cells after treatment with Hcy and (or) NaHS, bar: 25μm, n=3. (F) Schematic representation for detection of S-sulfhydration of target protein by maleimide assay. (G) S-sulfhydration level of Sp1 in mice fed with high Met diet for 8 weeks. (H) S-sulfhydration level of Sp1 in QSG-7701 cells after treatment with or without Hcy (2mM, 24h), n=5. (I) ChIP assay to determine relative Sp1 binding activity to the *Cse* promoter in QSG-7701 cells after treated with or without Hcy (2mM, 24h), n=3. (J) Reporter assay analysis of a series of deleted CSE promoter activity in QSG-7701 cells after treatment with or without Hcy (2mM, 24h), n=4. Graphs show means ± SEM. [Student’s t test, \**P*≤0.05, \*\*\**P*≤0.01, Met (Hcy) vs WT (Control)].

Given that Sp1 is the transcription factor of CSE in liver [33], we supposed that Sp1 might have happened to some changes. Firstly, we detected the protein level of Sp1 in the liver of mice and found no significant change (S1A Fig). Next, we treated QSG-7701 cells with Hcy for 24h, detected the protein level and sub-cellular location of Sp1. Similarly, no significant change was found (Fig 3E and S1C Fig). Then we turned our attention to the PTM of Sp1. Unlike CSE, Hcy had no significant influence on the nitration level of Sp1 either in vivo (S1D Fig) or in vitro (S1E Fig).

In view of H_2_S can S-sulfhydrate protein and affect its bio-activity, we wondered the impact of HHcy on S-sulfhydration and red maleimide was employed (S1F Fig). In WT mice and control group QSG-7701 cells (S1G Fig), we detected a large number of red bands, representing proteins with -SH as well as -SSH substitutes, with about 50% reduction after DTT treatment. However, less reduction was found in liver lysates of HHcy mice or QSG-7701 cells treated with Hcy (2mM, 36h). These results indicated the S-sulfhydration level of protein had been affected by Hcy both in vivo and in vitro.

It has been reported previously that Sp1 can be S-sulfhydrated by H_2_S [34, 35]. To substantiate the impact of Hcy on Sp1’s S-sulfhydration level, we detected the S-sulfhydration level of Sp1 by maleimide assay (Fig 3F, S1H Fig). As shown in Fig 3G, in the liver of mice with normal diet, the S-sulfhydration level of Sp1 was 58.74±12.28%, meanwhile in the high Met diet group, this value declined to 39.96±9.844%. What’s more, in normal QSG-7701 cells, the S-sulfhydration level of Sp1 was 40.16±4.018%, after treating with Hcy (2mM, 24h), it decreased to 15.24±3.492% (Fig 3H).

To further substantiate whether Hcy can affect the DNA binding activity of Sp1, ChIP assays were conducted. After treated with 2mM Hcy for 24h, the binding activity of Sp1 to *Cse*’s promoter declined about 24.5% (Fig 3I and S1I Fig).

Further, we utilized luciferase assay to determine whether Hcy regulated the activity of the *Cse* promoter. Six 5’-flanking fragments of *Cse* genomic promoters were amplified, sequenced and inserted into the upstream of firefly luciferase gene in the pGL3-Basic vector respectively, according to the published research [33] (S1 Table). Six vectors and the pGL3-Basic vector were respectively transfected into QSG-7701 cells together with Sp1 over-expression vector and pRL-TK vector transiently, and then the luciferase activity was assayed. The results showed that pCSE-185 (−44/+141) construct had no reporter activity, which was similar to the control vector pGL3-Basic. While pCSE-731 (−592/+139), pCSE-837(−698/+139), pCSE-937 (−794/+143) and pCSE-1335 (−1192/+143) fragments showed higher reporter activity. Additionally, pCSE-487 (−348/+139) showed less reporter activity (Fig 3J). In order to observe whether Hcy affects *Cse* promoter activity, the transfected cells were treated with 2mM Hcy for 24h. It was found that Hcy did inhibit the reporter activity of these fragments. These data further verified that Hcy could regulate *Cse* promoter activity on the transcriptional level.

The results mentioned above indicated that S-sulfhydration deficiency of Sp1 in HHcy could decrease its transcription activity, which was responsible for the decline of CSE’s transcription level.

### CSE was S-sulfhydrated at Cys84, Cys109, Cys172, Cys229, Cys252, Cys307 and Cys310

Similar to Sp1, CSE also has 10 cysteine residues which are the potential S-sulfhydrated sites [36]. In the present study, we used maleimide assay to detect the S-sulfhydration level of CSE (S2A Fig). As shown in Fig 4A, in the liver of WT mice, the S-sulfhydration level of CSE was 49.80±6.254%, while it declined to 21.50±10.81% in high Met diet group (17 weeks). Similarly, after treated with 2mM Hcy for 36h, the S-sulfhydrated level of CSE in QSG-7701 cells declined from 45.88±3.750% to 19.46±4.561% (Fig 4B).

**Fig 4.**
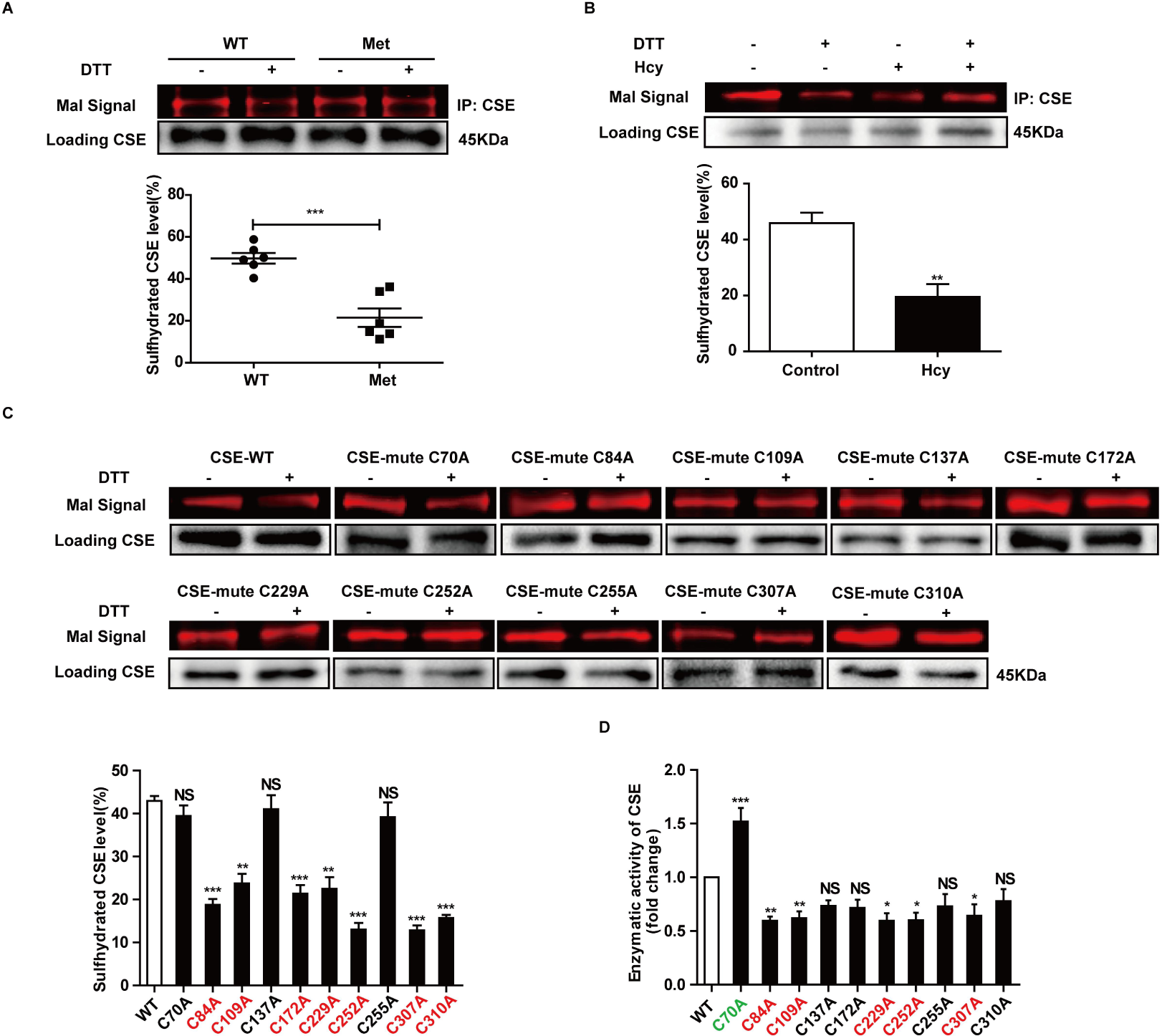
The S-sulfhydration site of CSE. (A) S-sulfhydration level of CSE in the liver of WT and HHcy mice. (B) S-sulfhydration level of CSE in QSG-7701 cells after treatment with or without Hcy (2mM, 36h), n=5. (C) CSE can be S-sulfhydrated on the residues Cys84, Cys109, Cys172, Cys229, Cys252, Cys307 and Cys310, n=3, 4. (D) Relative enzymatic activity of CSE expressed by different mutants compared to WT, n=4. Graphs show means ± SEM. [Student’s t test, (A, B) \**P*≤0.05, \*\*\**P*≤0.01, Met (Hcy) vs WT (Control), one-way ANOVA, (C, D), NS (no significant), \**P*≤0.05, \*\**P*≤0.005, \*\*\**P*≤0.001, Hcy (Met) vs Control (WT)].

To further determine the S-sulfhydrated site of CSE, we conducted site-directed mutation (S2 Table). Ten pEGFP-C3 plasmids with the mutation of cysteine residue in human CSE to alanine (C70A, C84A, C109A, C137A, C172A, C229A, C252A, C255A, C307A and C310A) together with the wild-type CSE (WT) plasmid were constructed and transiently transfected into HEK 293 cells respectively (S2B Fig). The physiological S-sulfhydration level of CSE was determined by maleimide assay (Fig 4C). Mutant C70A (39.45±4.211%), C137A (41.04±7.192%) and C255A (39.15±7.741%) showed a S-sulfhydration level similar to WT (42.92±1.976%), while others exhibited a significant decreased S-sulfhydration level, indicating that under physiological conditions, the S-sulfhydration of CSE occurred at Cys84 (18.74±2.341%), Cys109 (23.74±4.426%), Cys172 (21.35±4.432%), Cys229 (21.49±4.693%), Cys252 (23.03±3.029%), Cys307 (12.80±2.058%) and Cys310 (15.68±1.263%) residues.

Whether or not S-sulfhydration was necessary for the bio-activity of CSE? Next, we detected the enzymatic activity of CSE under physiological conditions. As shown in Fig 4D, mutant C84A, C109A, C229A, C252A and C307A exhibited a significant decrease of enzymatic activity, meanwhile there was no significant decrease in enzymatic activity of mutant C137A, C172A, C255A and C310A.

As mentioned above, these results indicated that S-sulfhydration was necessary for the function of CSE, and S-sulfhydration deficiency caused by Hcy could blunt the bio-activity of CSE.

### H_2_S donor rescued the bio-activity of Sp1-CSE-H_2_S pathway in HHcy

Next, we examined whether the supplement of H_2_S by its chemical donor (NaHS) could rescue the deficiency of CSE. After treated with NaHS, the level of H_2_S (Fig 5A) and CSE (Fig 5B, S3A Fig) had been restored. In vivo, we also found a similar phenomenon. In NaHS rescued group, the serum H_2_S (Fig 5C), tHcy (Fig 5D), 3-NT (S3B Fig), nitration (S3C Fig) and expression of CSE (Fig 5E, S3D Fig) were closed to WT. However, after treated with mithramycin A (MIA, Sp1’s inhibitor, Sigma ardrich) (S3E Fig) or siRNA of Sp1 (S3F Fig), the rescue ability of NaHS in QSG-7701 cells was abolished. These results indicated that the rescue ability of H_2_S donor was Sp1 dependent.

**Fig 5.**
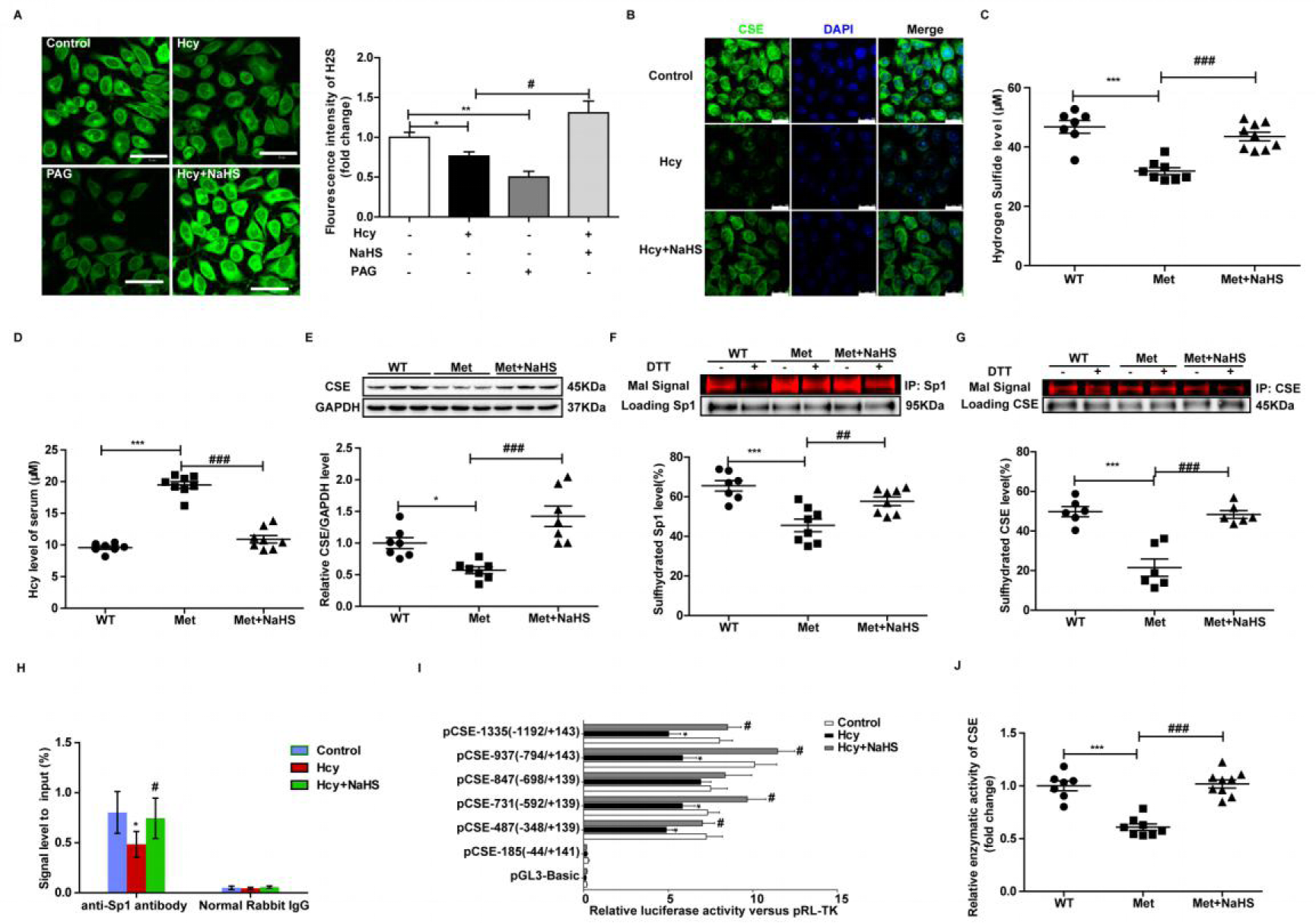
H_2_S donor rescued the bio-activity of Sp1-CSE-H_2_S pathway in HHcy. (A) H_2_S levels detected by C7-Az in QSG-7701 cells treated with Hcy (2mM), PAG (1mM, inhibitor of CSE’s enzymatic activity) and NaHS (2mM) for 36h, bar: 50μm, n=3. (B) NaHS treatment restored the level of CSE evidenced by immunofluorescence in vitro, n=3, bar: 25μm. (C) Serum H_2_S level of mice fed with high Met diet for 17 weeks. (D) Serum Hcy level of mice fed with high Met diet for 17 weeks. (E) The protein level of CSE in the liver of HHcy and NaHS rescued mice. (F) NaHS restored the S-sulfhydration level of Sp1 in HHcy mice. (G) NaHS restored the S-sulfhydration level of CSE in HHcy mice. (H) NaHS (2mM, 24h) restored the DNA binding activity of Sp1 in Hcy (2mM, 24h) treated QSG-7701cells, n=3. (I) NaHS (2mM, 24h) restored the promoter activity of *Cse* in Hcy (2mM, 24h) treated QSG-7701cells, n=4. (J) Relative enzymatic activity of CSE in the liver of HHcy mice. Graphs show means ± SEM. [one-way ANOVA, NS (no significant), \**P*≤0.05, \*\**P*≤0.01, \*\*\**P* ≤0.001, Hcy (Met) vs Control (WT), #*P*≤0.05, ##*P*≤0.01, ###*P* ≤0.001, Hcy (Met) + NaHS vs Hcy (Met)].

Further, we treated QSG-7701 cells with NaHS and found the S-sulfhydration deficiency of Sp1 and CSE in HHcy were restored to 32.96±5.019% and 42.08±5.944% respectively (S3G, H Fig). In vivo, both the S-sulfhydration level of Sp1 (57.69±6.249%) and CSE (48.34±4.833%) in the liver lysates of NaHS treated mice were nearly restored to WT mice (Fig 5F, G). Moreover, after treated with H_2_S donor, both the bio-activity of Sp1 (Fig 5H, I and S3I Fig) and CSE (Fig 5J) was nearly restored to the normal level.

These findings indicated that H_2_S could increase the S-sulfhydration level and restore the bio-activity of Sp1-CSE-H_2_S pathway, therefore block the progress of HHcy.

### S-sulfhydration deficiency followed the augment of nitration in HHcy

After treated QSG-7701 cells with Hcy (2mM, 24h), an increase of CSE’s nitration level was detected (Fig 1F). However, the S-sulfhydration level of CSE exhibited no significant change (Fig 6A). In addition, we detected the intracellular H_2_S in QSG-7701 cells and found an 18% decrease after 2mM Hcy treatment for 24h (Fig 2D). Further, we detected the S-sulfhydration level of Sp1 and found it decreased significantly (Fig 3H). In vivo, we identified the sequence of nitration increase and S-sulfhydration decline in mice fed with high Met diet, and a similar phenomenon was discovered (Fig 6B). Mice fed with high Met diet for 8 weeks exhibited an increase of tHcy (9.481±1.772μM) in serum (Fig 1A), along with an augmented 3-NT (Fig 1B) and nitrated CSE level in liver (Fig 1F). Meanwhile, a decline of Sp1’s S-sulfhydration was detected (Fig 3G). However, at that time, neither the S-sulfhydration of CSE (Fig 6C) nor the nitration of Sp1 (S1D Fig) exhibited any change. With the feeding continuing (to 17 weeks), both the tHcy (19.47±1.546μM) (Fig 1C), 3-NT (Fig 1D) and nitrated CSE (Fig 1G) level augmented further, accompanied by a further decline of H_2_S (Fig 6D). Then both the protein expression (Fig 3B) and S-sulfhydration deficiency of CSE (Fig 4A) were detected. Together, these results suggested that Hcy might increase the nitration level of CSE, led to a decline of its catalysate H_2_S. As a result, the S-sulfhydration level of Sp1 decreased and led to a decrease of CSE.

**Fig 6.**
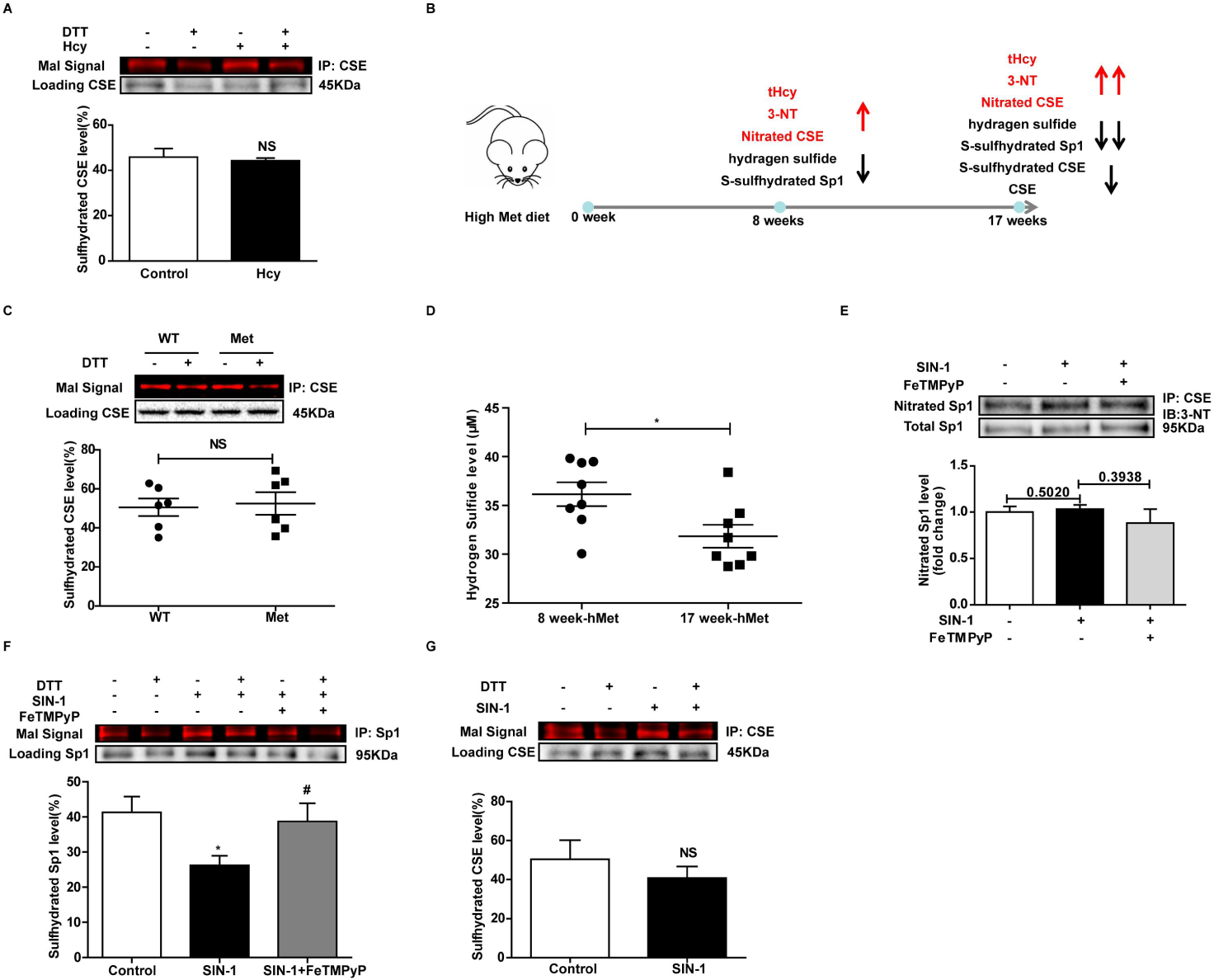
The S-sulfhydration deficiency followed the increase of nitration. **(A)** Treated QSG-7701 cells with Hcy (2mM, 24h), the S-sulfhydration level of CSE exhibited no significant change, n=3. **(B)** The sequence of nitration increase and S-sulfhydration decline in the progress of HHcy. **(C)** The S-sulfhydration level of CSE in the liver of mice fed with high Met diet for 8 weeks, n=6. **(D)** H_2_S level in the serum of mice, determined by methylene blue assay, n=8. **(E)** SIN-1 had no significant effect on the nitration level of Sp1 in QSG-7701 cells, n=4. **(F)** SIN-1 decreased the S-sulfhydration level of Sp1 in QSG-7701 cells, n=3. **(G)** SIN-1 had no significant effect on the S-sulfhydration level of Sp1 in QSG-7701 cells, n=3. Graphs show means ± SEM. [Student’s t test, (A, C, D, G), \**P*≤0.01. one-way ANOVA, (E, F), \**P*≤0.05, SIN-1 vs Control, #*P*≤0.05, SIN-1+FeTMPyP vs SIN-1].

To further confirm whether the S-sulfhydration deficiency was caused by the increased nitration level, SIN-1, a direct ONOO^-^ donor was used in vitro. As shown in Fig 2E, after treating with SIN-1, the nitration level of CSE augmented, sequentially decreased the intracellular H_2_S level (Fig 2F), meanwhile the nitration level of Sp1 remained unchanged (Fig 6E). Next, we found that SIN-1 decreased the S-sulfhydration level of Sp1 (Fig 6F). However, its effect on that of CSE showed no significance (Fig 6G). What’s more, by the use of FeTMPyP, the S-sulfhydration deficiency was partly rescued (Fig 6F).

As mentioned above, after augmenting the nitration level by SIN-1, the S-sulfhydration level of CSE decreased. In addition, blocking the increased ONOO^-^ by FeTMPyP reversed the S-sulfhydration deficiency. These results implied that the S-sulfhydration deficiency followed the augment of nitration level. And of note, blocking the increase of nitration might be a way to inhibit the decrease of S-sulfhydration in the early stage of HHcy.

## DISCUSSION

It has been indicated that HHcy induces nitration by up-regulating the ONOO^-^ level, which can blunt the bio-activity of proteins [37]. In the present study, we have discovered augmented 3-NT and nitration levels in HHcy model mice and Hcy-treated cells (see Figs 1B, 1D-1G), in keeping with our and others’ previous studies [12, 38]. The innovation of our present study is that CSE, one of the key enzymes for catalyzing the production of H_2_S in trans-sulfuration pathway, can be nitrated, and Hcy can augment its nitration level consequently blunting its enzymatic activity. This observation may explain a possible mechanism for H_2_S deficiency in HHcy.

Moreover, decreased expression levels of CSE have prompted us to examine a possible change in its transcription levels and we have found an S-sulfhydration deficiency of its transcription factor Sp1. It has been previously reported that Sp1 can be S-sulfhydrated by H_2_S. Saha et al [35] has found that S-sulfhydration of Cys68 and Cys755 residues by H_2_S stabilizes and enhances its binding to VEGFR-2 promoter, promoting the proliferation and migration of vascular endothelial cells. However, a study by Meng et al [34] have observed an opposite phenomenon that H_2_S S-sulfhydrates Cys664 residue of Sp1, blunting its binding activity with KLF5 promoter and resulting in the prevention of myocardial hypertrophy. Similar to other post-translational modifications, e.g. phosphorylation, under different pathological conditions, S-sulfhydration of Sp1 on different cysteine residues shows distinct impacts on its activity and exhibits different biological functions. Hence, to further illuminate the effect of S-sulfhydration on Sp1’s biological function in our model, ChIP assay has been used to measure binding activities of Sp1 to the promoter of *Cse* in QSG-7701 cells. We have found that S-sulfhydration deficiency of Sp1 is able to blunt its transcription activity, directly lowering the protein level of CSE. Thus, the S-sulfhydration of Sp1 is necessary for the transcription of *Cse* in Hcy-metabolism.

Furthermore, we have also observed that CSE can be S-sulfhydrated and Hcy can affect its S-sulfhydration levels as well. Next, by site-directed mutation assay, we have detected 7 possible S-sulfhydration sites in CSE under the physiological condition. In theory, S-sulfhydration levels of CSE should decrease around 14% after mutating each site, but our results have shown a reduction of almost 50%. This disproportionate decline has also been observed by others. For example, Ohno et al [39] have demonstrated that both Cys71 and Cys124 in PTEN are targets for S-sulfhydration by site-directed mutation. In their study, when singly mutating Cys71, the S-sulfhydration of PTEN disappears, a phenomenon similar to the co-mutation of Cys71 and Cys124. However, when mutating Cys124 alone, the S-sulfhydration level of PTEN decreases nearly 90%. In another report, Zheng et al [40] have found that RUNX2 can be S-sulfhydrated in Cys123 and Cys132 residues, and when singly mutating Cys123 or Cys132, the S-sulfhydration level of RUNX2 decreases nearly 90%. Coincidently, Du et al [41] have also observed a similar phenomenon in which they have found that Sirt1 can be S-sulfhydrated at its two zinc finger domains, while mutating anyone of the two domains, S-sulfhydration of Sirt1 can be completely abolished. Thus, the number of mutation sites is not strictly correlated with S-sulfhydration levels, suggesting a synergistic effect or complementary role from the remaining amino acid residues in the protein. Strictly speaking, the present study is a rather preliminary discussion about CSE’s S-sulfhydration sites under the physiological condition. It may be insufficient to determine the modified sites by site-directed mutation, and to be precise, the S-sulfhydrated sites need to be identified by a combined detection employing site-directed mutation and Mass spectrometry analysis (MS) in the future. It is worth noting that a recent report has studied the S-sulfhydration of CSE in the metabolism of Hcy [25], which has shown similar results as ours in that HHcy decreases the S-sulfhydration level of CSE in mice. The differences are that, when mutating four cysteine residues (Cys252, Cys255, Cys307, Cys310) located in the C-terminal of CSE, all these four cysteine residues could be S-sulfhydrated under the physiological condition [25]. In our study, however, Cys255 is not able to be S-sulfhydrated (see Fig 4C). This distinction might be caused by the detection methods used for S-sulfhydration as Fan et al. used a modified biotin switch assay [25] but we employed a maleimide assay (see Methods section and Fig 3F). As maleimide selectively interacts with sulfhydryl groups of cysteines, the maleimide assay by labeling both S-sulfhydrated as well as unsulfhydrated cysteines is advantageous in that nitrosylated or oxidized cysteines do not react with maleimide [42], suggesting that the maleimide assay we used may be more accurate in detecting S-sulfhydration.

By uncovering the effect of S-sulfhydration on CSE’s enzymatic activity, we have found some mutations that inhibit the bio-activity of CSE. The S-sulfhydration is functional on some sites (Cys84, Cys109, Cys229, Cys252, and Cys307) but not on other sites (Cys172 and Cys310). Interestingly, cysteine residue Cys70 seems to be distinctive although its mutation to alanine does not affect the S-sulfhydration level of CSE with increased enzymatic activity. This distinctive result suggests that Cys70 appears to be another kind of PTM, e.g., S-nitrosylation, which always occurs at the cysteine residue and blunts the function of protein[36]. Therefore, it may be of significance to focus on CSE’s S-nitrosylation in future studies. Metabolic regulations in organisms are extremely complex and always inter-related each other, especially in the PTM of functional proteins. Sen et al. [42] have established that TNF-α treatment firstly induces the S-sulfhydration of NF-κB’s large subunit P65, enhances its binding to Ribosomal Protein S3. However, when treating time extends, S-sulfhydration is replaced by S-nitrosylation. Feng et al. [43] have reported that NO-induced ERK S-nitrosylation inhibits its phosphorylation and triggers apoptotic program. Altaany et al. [44] have revealed that H_2_S S-sulfhydrated eNOS increases its activity, promotes its phosphorylation and inhibits its S-nitrosylation. In the present study, we have detected that the augmentation of nitration induced by Hcy can cause a deficiency of S-sulfhydration through the mediation of CSE and H_2_S. In addition, we have also observed that Hcy induces S-sulfhydration deficiency of Sp1 prior to that of CSE. Under the same treatment, S-sulfhydration levels of CSE and Sp1 induced by H_2_S deficiency are different, prompting us to examine the underlying mechanism(s). As shown in Fig 2D, we utilize H_2_S-selective sensor C7-Az to detect intracellular H_2_S and have found primary existence of H_2_S in the cytoplasm of QSG-7701 cells. The level of H_2_S in nucleus is rather low compared with that in the cytoplasm, which is consistent with the subcellular localization of CSE (Fig 3C). These findings suggest that, under the physiological conditions, the basal level of H_2_S in the cytoplasm is much higher than that in the nucleus. Thus, the fluctuation of H_2_S in the nucleus induced by HHcy may be more dramatic than that in the cytoplasm. In other words, the nucleoproteins may be more sensitive to the decrease of H_2_S. Hence, the S-sulfhydration of Sp1 decreases after treating with Hcy while the S-sulfhydration of CSE remains almost unchanged. With HHcy aggravating, the level of H_2_S decreases further, destroys the anti-fluctuation ability of H_2_S in the cytoplasm and finally, causes the S-sulfhydration deficiency of plasmosin. By this time, the S-sulfhydration deficiency of CSE has occurred.

HHcy can be induced by many factors, among which high methionine diet plays an important role. The metabolism of Hcy is at the intersection of two metabolic pathways: remethylation and trans-sulfuration. Previous studies have demonstrated that the utilization of homocysteine molecules by the trans-sulfuration and remethylation pathways is nutritionally regulated [45, 46]. When excess dietary methionine is administered, remethylation pathway shows to be inhibited and the trans-sulfuration pathway acts as the protagonist [19]. In order to rule out other factors that may also cause HHcy, we fed the mice with high Met diet to establish HHcy model. Hence, it is rational to focus on the trans-sulfuration pathway primarily. Through the work presented here, we have confirmed that the augmented nitration levels induced by Hcy lead to an S-sulfhydration deficiency of Sp1 and CSE, which consequently impairs the trans-sulfuration pathway. However, under such circumstances, it would be interesting to investigate if the remethylation pathway could be restarted or if it could be regulated by H_2_S, which is a product of the trans-sulfuration pathway.

After the mice fed with high Met diet for 8 weeks, tHcy levels in the serum were increased to 9.481±0.06265 μM, significantly higher than those in WT (6.607±0.3299 μM). In the mice fed for 17 weeks, tHcy levels in the serum were increased from 9.581±0.7323 μM (WT) to 19.47±1.546 μM (Met group). We had not found any criterion that could estimate the level of tHcy in mice (above 15 μM for humans). Therefore, the 8 week-fed mice should not be categorized into HHcy group whom we termed as pre-HHcy mice in order to differentiate them from HHcy mice (17 week-fed mice). In the pre-HHcy mice, we found an augmented nitration level and, with time elapsing, the augmented nitration level led to S-sulfhydration deficiency. Meanwhile, S-sulfhydration deficiency weakened the bio-activity of Sp1-CSE-H_2_S pathway which, in turn, blunted the enzyme’s ability to metabolize Hcy. By this time, the metabolism of Hcy was falling into a vicious circle.

A wide range (from 0.1 μM to more than 300 μM) of circulating H_2_S levels has been reported under the physiological conditions. This remarkable variation is largely attributable to the wide array of measuring methods used [13]. In the present study, we have selected methylene blue assay, a classic method universally used to determine H_2_S in vivo. The decline of H_2_S is not strictly consistent with the decreases S-sulfhydration, which might be caused by the low sensitivity of the detection method. This is a limitation of this study in terms of determining precise levels of H_2_S and other methods with higher sensitivities should be trialed in the future.

In conclusion, we have, for the first time, revealed a possible underlying mechanism involved in the inactivition of Sp1-CSE-H_2_S pathway in hyperhomocysteinemia development. It is worth noting that we have also uncovered an important role of PTM in the progress of HHcy (Fig 7). Accumulation of Hcy augments the nitration of CSE and lowers H_2_S production, which subsequently decreases the S-sulfhydration of Sp1. Insufficient S-sulfhydration of Sp1 diminishes the transcription of *Cse*. The level of H_2_S decreases further with lapsed time, which, in turn, induces the S-sulfhydration deficiency of CSE. In the end, declines in both quality and quantity of CSE may occur, leading to a further increase in serum Hcy, and as a result, the progress of HHcy falls into a vicious circle. This study provides a novel insight into the progress of HHcy in terms of the mechanism, which offers a potential therapeutic strategy for HHcy through targeting the PTM of Sp1-CSE-H_2_S pathway.

**Fig 7.**
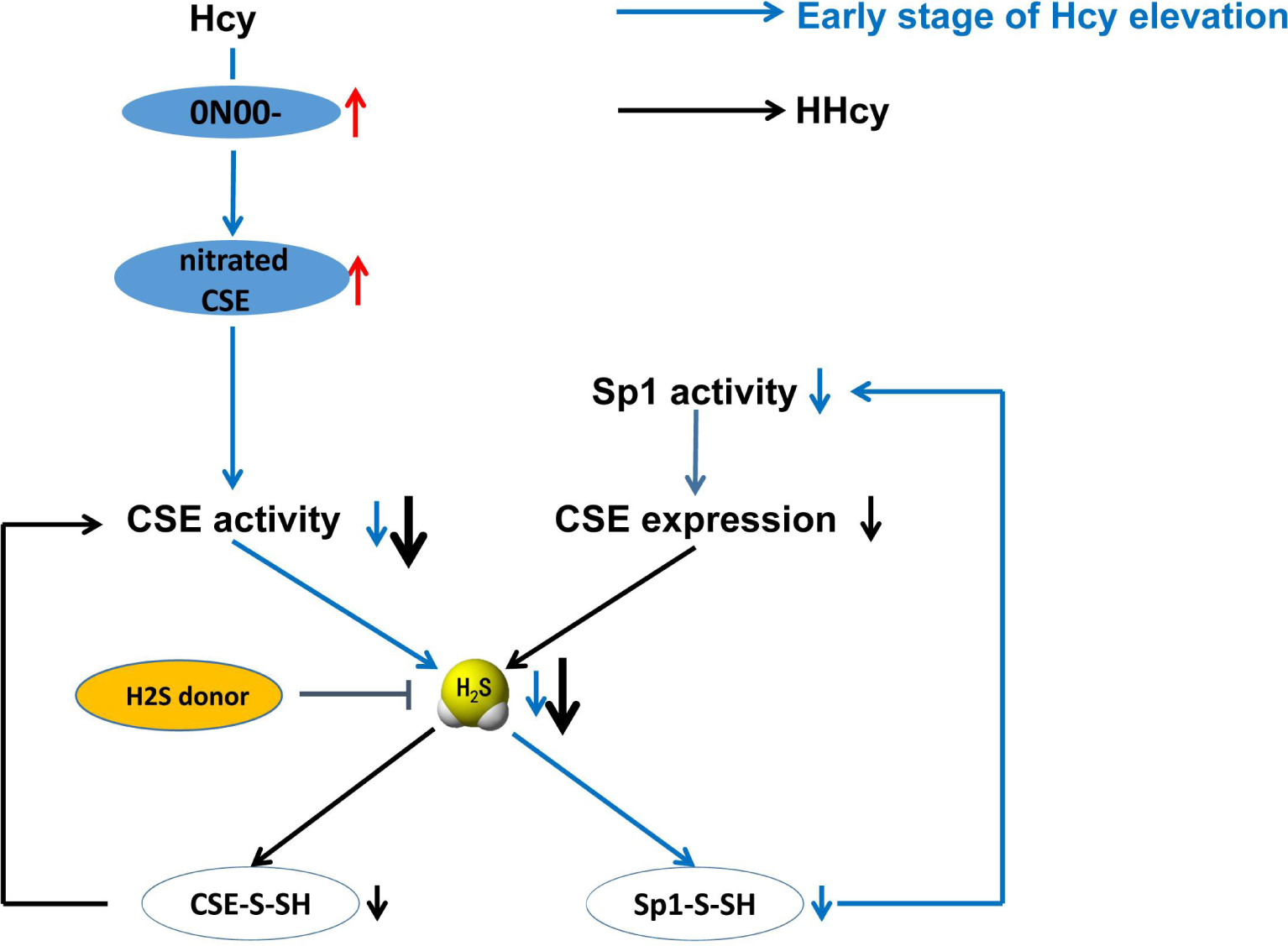
Schematic diagram of the present findings. At the early stage of Hcy elevation, ONOO^-^ level increased significantly, which caused the augment of nitrated CSE and resulted in the reduction of its catalyst H_2_S. The decrease of H_2_S level caused the S-sulfhydration deficiency of Sp1. S-sulfhydration of Sp1 was necessary for its transcription activity, consequently, the decline of *Cse*’s mRNA level happened, and then resulted in the decrease of CSE. With time continuing, the CSE deficiency lowered the production of H_2_S further, which finally led to the S-sulfhydration deficiency of CSE. Because of their combined impact, the expression and enzymatic activity of CSE decreased further, the progress of HHcy may develop into a vicious cycle. Of note, H_2_S donor could rescue the activity and expression level of Sp1-CSE-H_2_S pathway by increasing the S-sulfhydration level.

## MATERIALS AND METHODS

### Reagents and antibodies

Homocysteine (Hcy), sodium hydrosulfide (NaHS), mithramycin A (MIA), dithiothreitol (DTT), L-cysteine (Cys) and pyridoxal 5’-phosphate (PLP) were purchased from Sigma-Aldrich. Alexa Fluor 680 conjugated C2 maleimide was purchased from Invitrogen. The following antibodies were used in this work: anti-CSE antibody (Proteintech, 12217-1-AP), anti-Sp1 antibody (Santa Cruz, sc-17824), anti-Sp1 antibody (Cell Signaling Technology, D4C3), anti-GAPDH antibody (Sigma-Aldrich, G9295), anti-β-tublin antibody (ABclonal, AC031), anti-Lamin B1 antibody (Proteintech, 66095-1-lg), anti-Nitrotyrosine antibody (Merck Millipore, 05-233), non-specific mouse IgG (Beyotime, A7028), Normal Rabbit IgG (Cell Signaling Technology, 2729).

### Animals and cell culture

Healthy male C57BL/6J mice (8-week-old, SPF grade, n=10 each group) were supplied by the Animal Center, Capital Medical University. HHcy models were induced by feeding mice with high methionine diet (2.5% Met diet) for 17 weeks. The NaHS rescue group mice were intraperitoneal injected with NaHS (diluted in normal saline, 10mg/Kg) every other day. Mice fed on common diet were used as control (WT). All animals received human care in compliance with institutional guideline and the “Guide for the Care and Use of Laboratory Animals” prepared by the “Institute of Laboratory Animal Resources, Commission on Life Sciences, National Research Council”.

QSG-7701 cells were used for in vitro research and were maintained at 37℃ with 5% CO_2_ in DMEM/HIGH GLUCOSE media (Hyclone, Logan, UT) containing phenol red supplemented with 10% fetal calf serum and 100U/mL penicillin/1000μg/mL streptomycin.

### Measurement of Hcy and H_2_S level

After euthanized the mice, blood samples were collected from carotid arterial cannulation containing 7.5% EDTA-Na_2_ (15ml/L), and immediately centrifuged at 3000g, 4°C for 10 min. Serum was separated and measured the Hcy and H_2_S level.

### CSE activity assay

CSE activity assays were performed using cysteine, as substrates, as reported earlier [42]. Briefly, cells were homogenized in PBS (pH 6.8) containing 13 protease inhibitor cocktail (1mM, final concentration). The enzymatic assay was performed in 250μl reaction mixture containing the cell homogenate (equal protein from cells), PBS (pH 7.4), 10mM cysteine and 15μM PLP. The reaction mixture was flushed with nitrogen and sealed, and the reactions were initiated by transferring the vials from ice to a 37℃ incubator shaker. After incubating for 60min, the H_2_S level was measured with hydrogen sulfide detection kit (Jiancheng Bioengineering Institute, Nanjing, China).

### Immunoprecipitation analysis

For immunoprecipitation assay, QSG-7701 cells were homogenized in lysis buffer (Tris-HCl 20mM, Triton 0.1%, NaCl 100mM, PMSF 100μM), then centrifuged at 12,000rpm for 15min at 4°C. Cell lysates (500μg total protein) were incubated with 20μl Protein A/G PLUS-Agarose (Santa Cruz) by occasional gentle mixing for 6h at 4°C, then centrifuged at 2500g for 3min at 4°C. The supernatant was incubated with 1μg anti-CSE antibody (Proteintech) or 2μg anti-Sp1 antibody (Santa Cruz) with occasional gentle mixing for 8h or overnight at 4°C, then added 20μl Protein A/G PLUS-Agarose (Santa Cruz) and co-incubated for 4h. Beads were pelleted by centrifuging at 2500g for 3min at 4°C, washed with washing buffer (Tris-HCl 20mM, Triton 0.1%, NaCl 300mM) 3 times for 10 min each and finally added with 20μl 2×SDS-PAGE buffer and boiled at 99°C for 10min followed by gel electrophoresis. Gels were transferred to PVDF membrane (Millipore). The membranes were blocked with 5% nonfat-dry milk in Tris-buffered saline containing 0.1% Tween (TBST, pH 7.6). The primary antibodies used for immunoblotting were anti-Nitrotyrosine antibody (1:1000, Millipore). Membranes were then washed with TBST (3 times for 10min each), incubated with secondary antibodies coupled to horseradish peroxidase, washed again with TBST (3 times for 10min each). ECL Plus substrate (Millipore) was applied to the imaging, and images were captured in a gel documentation system (Bio-Rad). Relative optical density of protein bands was analyzed using gel software Image Lab 3.0. Then regenerated the member with Western Stripper (Pulilai, Beijing, China) and washed it with TBST then detected loading protein with anti-CSE antibody (1:1000, Proteintech) or anti-Sp1 antibody (1:1000, Santa Cruz).

### Measurement of H_2_S level by an H_2_S-selective sensor

The specific fluorescent probe C-7Az was kindly gifted by Dr. Xinjing Tang (Peking University). The H_2_S level in QSG-7701 cells was measured using C-7Az as previously described [47]. In brief, the cultured cells were washed with PBS 3 times, then 20μM (final concentration) C-7Az dissolved in PBS was added and incubated at 37°C for 30min, followed by washing of the cells with PBS buffer. Cell imaging was carried out by two-photon con-focal laser scanning fluorescence microscopy with the excitation of a 720nm laser.

### Maleimide assay

This assay was designed based on the principles described before [42]. With some modifications, briefly, QSG-7701 cells were homogenized in lysis buffer (Tris-HCl 20mM, Triton 0.1%, NaCl 100mM, PMSF 100μM), then centrifuged at 14,000rpm for 15min at 4°C. Cell lysates (300μg total protein) were incubated with 20μl Protein A/G PLUS-Agarose (Santa Cruz) by occasional gentle mixing for 12h at 4°C to remove the non-specific binding proteins, and then centrifuged at 2500g for 3min at 4°C. The supernatant was removed into another clear tube and incubated with 1μg anti-CSE antibody (Proteintech) or 2μg anti-Sp1 antibody (Santa Cruz) with occasional gentle mixing for 8h at 4°C, 1μg appropriate control IgG was added into another group. Then added 40μl Protein A/G PLUS-Agarose (Santa Cruz) and co-incubated for 8h. Beads were pelleted by centrifuging at 2500g for 3min at 4°C, washed with lysis buffer once for 1h, and then followed by washing buffer (Tris-HCl 20mM, Triton 0.1%, NaCl 500mM) once for 15min. Next incubated with Alexa Fluor 680 conjugated C2 maleimide (red maleimide, 1mM, final concentration) and kept for 2h at 4°C with occasional gentle mixing. Beads were pelleted by centrifuging at 2500g for 3min at 4°C, washed with lysis buffer once for 1h, and then followed by washing buffer once for 15min. Then re-suspended the beads with 1ml lysis buffer and divided equally into two tubes, among which one treated with DTT (1mM, final concentration), another without DTT, both incubated for 1h at 4°C with occasional gentle mixing. Beads were pelleted and washed with lysis buffer once for 1h, and then followed by washing buffer once for 15min. Lastly, Beads were precipitated by centrifuging at 2500g for 3min at 4°C, added with 20μl 2×SDS-PAGE buffer, and then boiled at 99°C for 10min followed by gel electrophoresis. The gel was scanned with the Li-COR Odyssey system (Li-Cor Biosciences, Lincoln, NE, USA), the intensity of red fluorescence of CSE or Sp1 was quantified using software attached to the Odyssey system. These membranes were next employed for western blotting with anti-CSE antibody or anti-Sp1 antibody to detect the loaded CSE or Sp1.

### Immunofluorescence staining

Cells were washed with pre-cooling PBS three times, fixed with 10% triformol for 30min, then blocked with 5% BSA (dissolved in PBS) for 30min. Incubated with anti-CSE antibody (1:200, Proteintech) or anti-Sp1 antibody (1:200, Proteintech) overnight. After additional washing, cells were incubated with directly conjugated fluorescent secondary antibodies and DAPI (Invitrogen). Fluorescence was imaged using a confocal microscopy. The intensity of fluorescence was quantified using the histogram function in NIH Image J.

### Modified maleimide assay to detect overall protein S-sulfhydration

20μg total protein was added with 5μl 5×SDS-loading buffer, kept the final volume at 25μl, then incubated with Alexa Fluor 680 conjugated C2 maleimide (red maleimide, 1mM, final concentration) and kept for 2h at 4°C with occasional gentle mixing. Then DTT (1mM, final concentration) was added to the DTT treated group and incubated at 4°C for 1h. The reaction was terminated by adding methanol-chloroform-H_2_O and precipitated proteins were dissolved in 20μl lysis buffer (Tris-HCl 20mM, Triton 0.1%, NaCl 100mM, PMSF 100μM). Lastly, added 5μl 5×SDS-loading buffer into the sample and boiled at 99°C for 10min followed by gel electrophoresis. The gel was scanned by Li-COR Odyssey system (setting 0.5; Li-Cor Biosciences, Lincoln, NE, USA) to detect and qualify the red fluorescence of protein bands. Then, loaded protein bands were detected by Coomassie brilliant blue staining.

### Sp1 siRNA transfection

siRNA transfection of QSG-7701 cells in 6-well plate with Sp1siRNA (Santa Cruz, sc-29487) was carried out using Lipofectamine**^®^** RNAiMAX Reagent. Briefly, cells at 30-40% confluence were plated in 6-well plate and grown overnight in complete DMEM/HIGH GLUCOSE media (Hyclone, Logan, UT). Before transfection, cells were washed once with PBS, then added 1ml DMEM medium without antibiotic into each well and kept in CO_2_ incubator until addition of transfection mixture. 60pmol Sp1siRNA and 9μl Lipofectamine**^®^** RNAiMAX Reagent (Invitrogen) diluted with 150μl Opti-MEM**^®^** Medium (Gibico) respectively were prepared for each well. After placing at room temperature for 5min, the diluted reagent were mixed and then placed at room temperature for another 5min. Next, 150μl mixture was added into each well and kept in CO_2_ incubator. After 8h, NaHS was added to the plates. The cells were cultured for 36h before any subsequent analysis.

### Chromatin immunoprecipitation

Chromatin immunoprecipitation (ChIP) assays were performed using #9004 Simple ChIP Plus Kit (Agarose Beads) (CST) as manufacturer’s instructions. Briefly, cells were treated with 1% (final concentration) formaldehyde to cross-link proteins to DNA. After cross-linking, cells were lysed and sonicated to shear the chromatin to a manageable size (200-1000bp). Immunoselections of cross-linked protein-DNA were performed with anti-Sp1 antibody (D4C3, CST) or normal rabbit IgG #2729 (negative control), and ChIP grade protein G-conjugated agarose beads. Protein-DNA complexes were washed, and then protein-DNA cross-links were reversed to free DNAs. The quantitative analysis of Sp1 and the *hCSE* gene interaction was determined by conventional PCR and real-time PCR using primers for *CSE* promoter as follows. *hCSE*: forward, 5’-GGACTCCAGCTTCACTCCGCTT-3’; reverse, 5’-TTAGCGGGTCTGCAGTCTCACG-3’。

### Luciferase assay

Cells were plated per well of a 24-well plate and grown overnight. On the day of transfection, the cells were washed once with PBS, and then 200μl of DMEM medium without antibiotic was added to each well before adding transfection mixture. Transfection mixture per well was prepared with 3μl Lipo6000^TM^ transfection reagent (C0526, Beyotime, Shanghai, China), 500ng promoter-firefly plasmid, 500ng Sp1 plasmid and 500ng wild-type pRLTK-Renilla plasmid diluted in 150μl Opti-MEM**^®^** Medium following the manufacturer’s protocol. After addition of transfection mixture, cells were kept in a CO_2_ incubator for 6h then replaced the medium with 500μl complete DMEM medium with or without Hcy (and or NaHS). Luciferase activity was measured using a Dual Luciferase Reporter Assay Kit (Vazyme, Nanjing, China) following the suppliers’ protocol 36h post transfection.

### Site directed mutation and cell transfection

The mutation of cysteine residue in human CSE to alanine in pEGFP-C3 plasmids were generated commercially by LKL Company (Beijing, China). HEK 293 cells were dissociated with trypsin, quantified and plated in 10cm plates. The next day, when the cell density up to 80%, transfected with 15μg of each vector using Lipo6000^TM^ reagent (Beyotime, Shanghai, China) and Opti-MEM^®^ Medium (Gibico) according to the manufacturer’s instructions. Six hours after transfection, the transfected medium was replaced with complete DMEM/HIGH GLUCOSE media (Hyclone, Logan, UT). Cultured for another 48h, then the cells were harvested with 200μl lysis buffer (Tris-HCl 20mM, Triton 0.1%, NaCl 100mM, and PMSF 100μM). 150μg total protein was used for maleimide assay to detect the S-sulfhydration level of CSE.

### Statistical analyses

Data were represented as the means ± SEM. Unpaired Student’s t test was used to analyze data with only two sets. One-way analysis of variance (ANOVA) was performed to determine whether there was a significant difference between more than two data sets, followed by Bonferroni’s post hoc test, using Graph Pad Prism 5.0. Group differences at the level of *P*< 0.05 were considered statistically significant. Asterisk (*) represented *P*< 0.05; double asterisk (**) represented *P*< 0.01; triple asterisk (***) represented *P*< 0.001.

## ACKKNOWLEDGMENTS

We thank Dr. Bin Geng and Dr. Xinjing Tang from School of Medicine, Peking University, Beijing, China, for help in the detection of H_2_S in living cells; we are grateful to Dr. Wenjie Zhang from School of Medicine, Shihezi University, Xinjiang, China, for assistance in review and language editing.

## Author contributions

Wen Wang designed the experiments; Chenghua Luo performed most of the experimental analysis and wrote the manuscript; Dengyu Ji, Yan Li, Yan Cao, Shangyue Zhang, Wenjing Yan, Ke Xue and Jiayin Chai performed some of the experiments; Ye Wu and Huirong Liu provided technical support. All authors commented on and approved the manuscript.

---

**S1 Fig.**
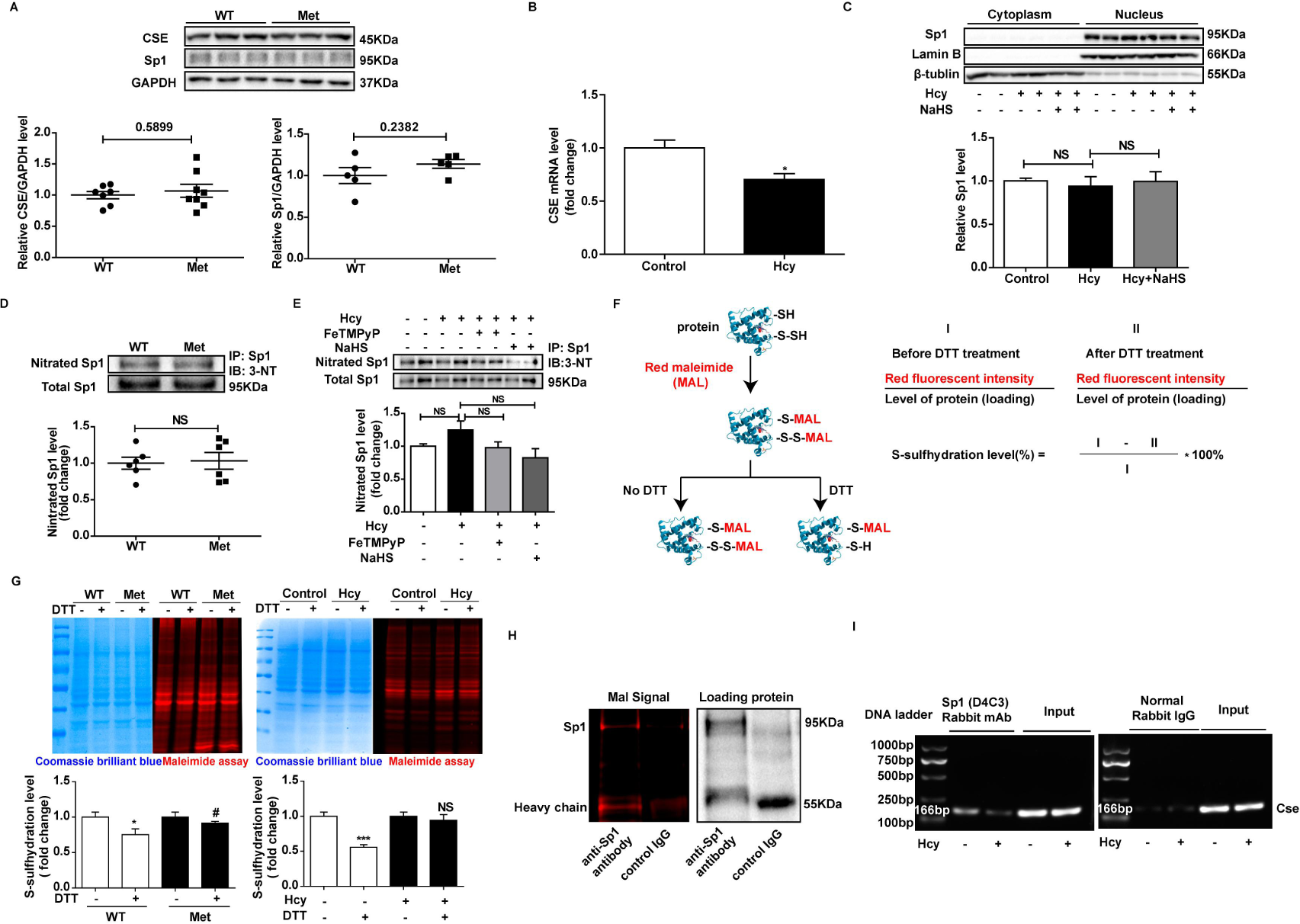
The effect of Hcy on S-sulfhydration. (A) Protein level of CSE and Sp1 in the liver of mice fed with high Met diet for 8 weeks. (B) Relative mRNA abundance of Cse in QSG-7701 cells determined by qRT-PCR, cells were treated with or without 2mM Hcy for 24h, n=3. (C) Protein level and subcellular location of Sp1 in QSG-7701 cells after treatment with Hcy and (or) NaHS, n=3. (D) Nitration level of Sp1 in the liver of mice fed with high Met diet for 8 weeks. (E) Nitration level of Sp1 in the liver of QSG-7701 cells, n=3. (F) Schematic representation for detection of S-sulfhydration of over-all protein by maleimide assay. (G) Red maleimide assay was performed to detect the S-sulfhydration of total protein in liver tissue lysates of both wild type and mice fed with high Met diet for 17 weeks, n=6. Red maleimide assay was performed to detect the S-sulfhydration of total protein in QSG-7701 cells treated with or without Hcy (2mM, 36h, n=3). (H) Negative control for maleimide assay of Sp1. (I) ChIP assay to determine relative Sp1 binding activity to the Cse promoter in QSG-7701 cells after treated with or without Hcy (2mM, 24h), the DNA amplification of Cse gene by conventional PCR, n=3. Graphs show means ± SEM. [Student’s t test, (A, B, C, D, E, G), NS (no significant), *P≤0.05, **P≤0.01, Met (Hcy) vs WT (Control). one-way ANOVA, (D, F), NS (no significant), *P≤0.05, ***P≤0.001, Hcy (Met) vs Control (WT). #P≤0.05, Hcy+FeTMPyP vs Hcy]. Cells were treated with Hcy (2mM), NaHS (2mM), FeTMPyP (100μM) for 24h then harvested for detection.

**S2 Fig.**
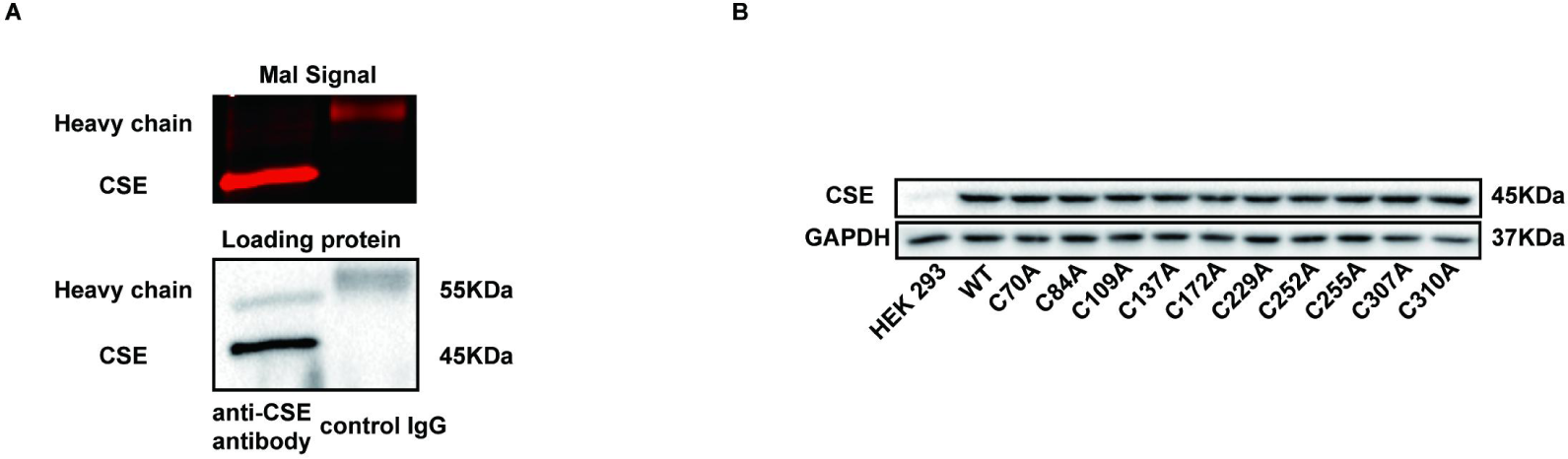
The effect of Hcy on S-sulfhydration. (A) Negative control for maleimide assay of CSE. (B) The expression of CSE in HEK 293 cells was determined by Western Blot.

**S3 Fig.**
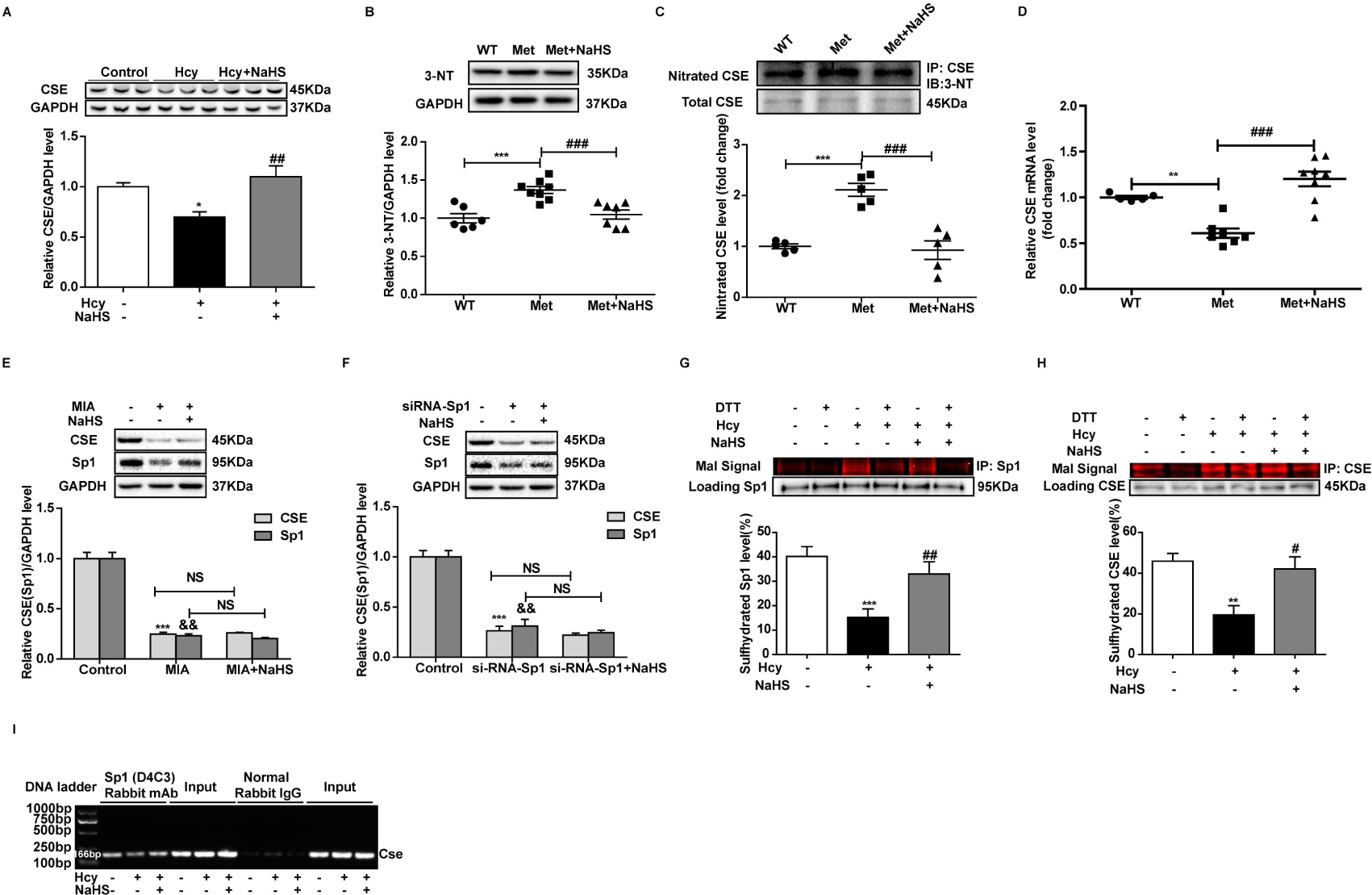
Hydrogen sulfide donor rescued the progress of HHcy. (A) NaHS treatment restored the level of CSE evidenced by immunoblot assay in vitro, n=3. (B) 3-NT level in liver lysate of mice fed with high Met diet for 17 weeks. (C) The nitration level of CSE in liver lysate of mice fed with high Met diet for 17 weeks. (D) The mRNA level of *Cse* in liver of HHcy and NaHS rescued mice. (E) Preconditioned the QSG-7701 cells with Sp1’s specific inhibitor mithramycin A (MIA, 0.5μM, 36h), n=4. (F) Knockdowned Sp1 by siRNA in the QSG-7701 cells, NaHS failed to restore the CSE level, n=4. (G) The H_2_S donor NaHS (2mM, 24h) restored the S-sulfhydration level of Sp1 in Hcy (2mM, 24h) treated QSG-7701cells, n=4. (H) The H_2_S donor NaHS (2mM, 24h) restored the S-sulfhydration level of CSE in Hcy (2mM, 36h) treated QSG-7701cells, n=4. (I) ChIP assay to determine relative Sp1 binding activity to the *Cse* promoter in QSG-7701 cells, the DNA amplification of *Cse* gene by conventional PCR, n=3. Graphs show means ± SEM. [one-way ANOVA,\**P*≤0.05, \*\**P*≤0.01, \*\*\**P* ≤0.001, Hcy (Met) vs Control (WT), #*P*≤0.05, ##*P*≤0.01, ###*P* ≤ 0.001, Hcy (Met) + NaHS vs Hcy (Met)].

**S1 Table.**
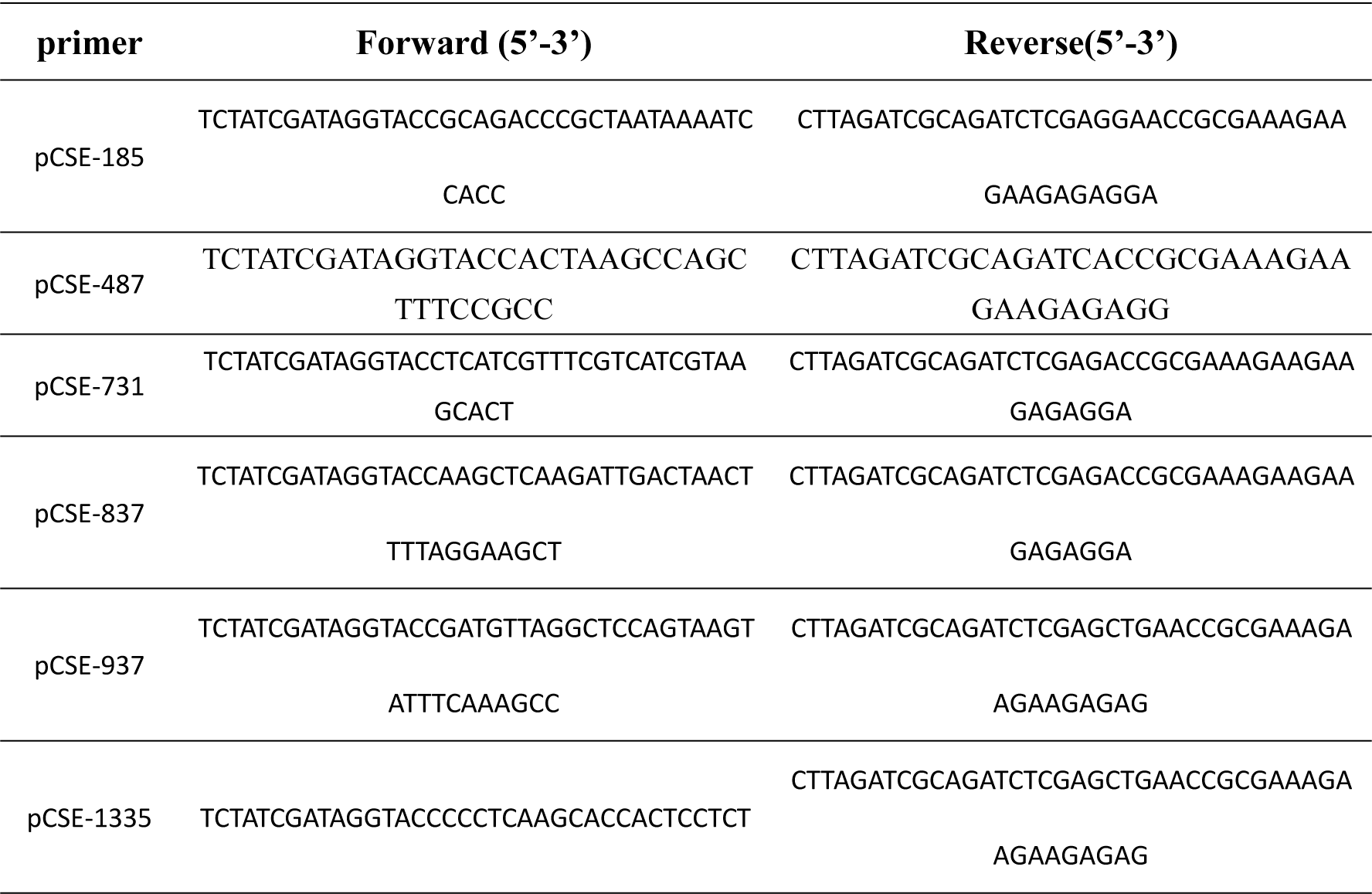
Primers for the construction of vectors used in luciferase assay.

**S2 Table.**
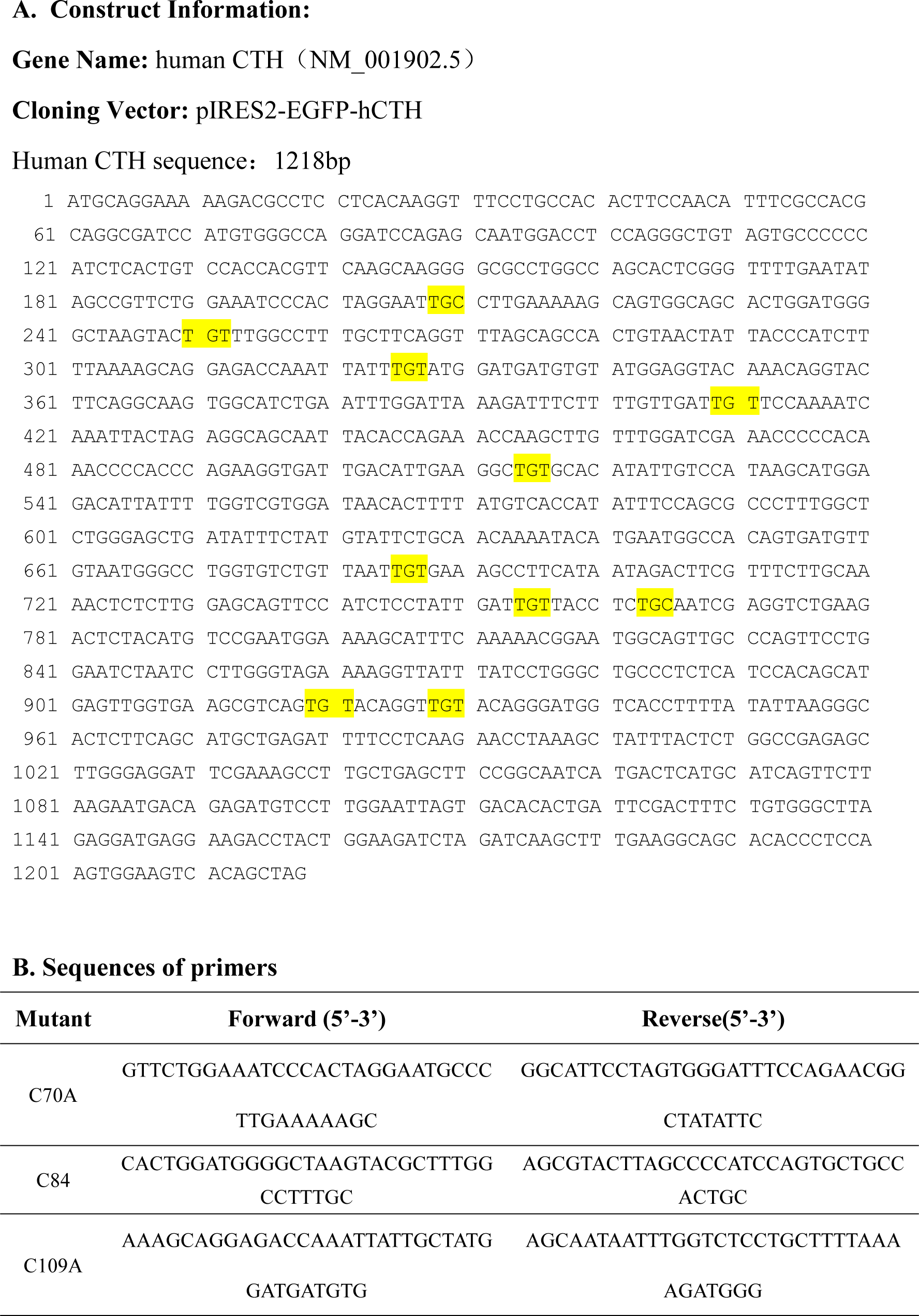
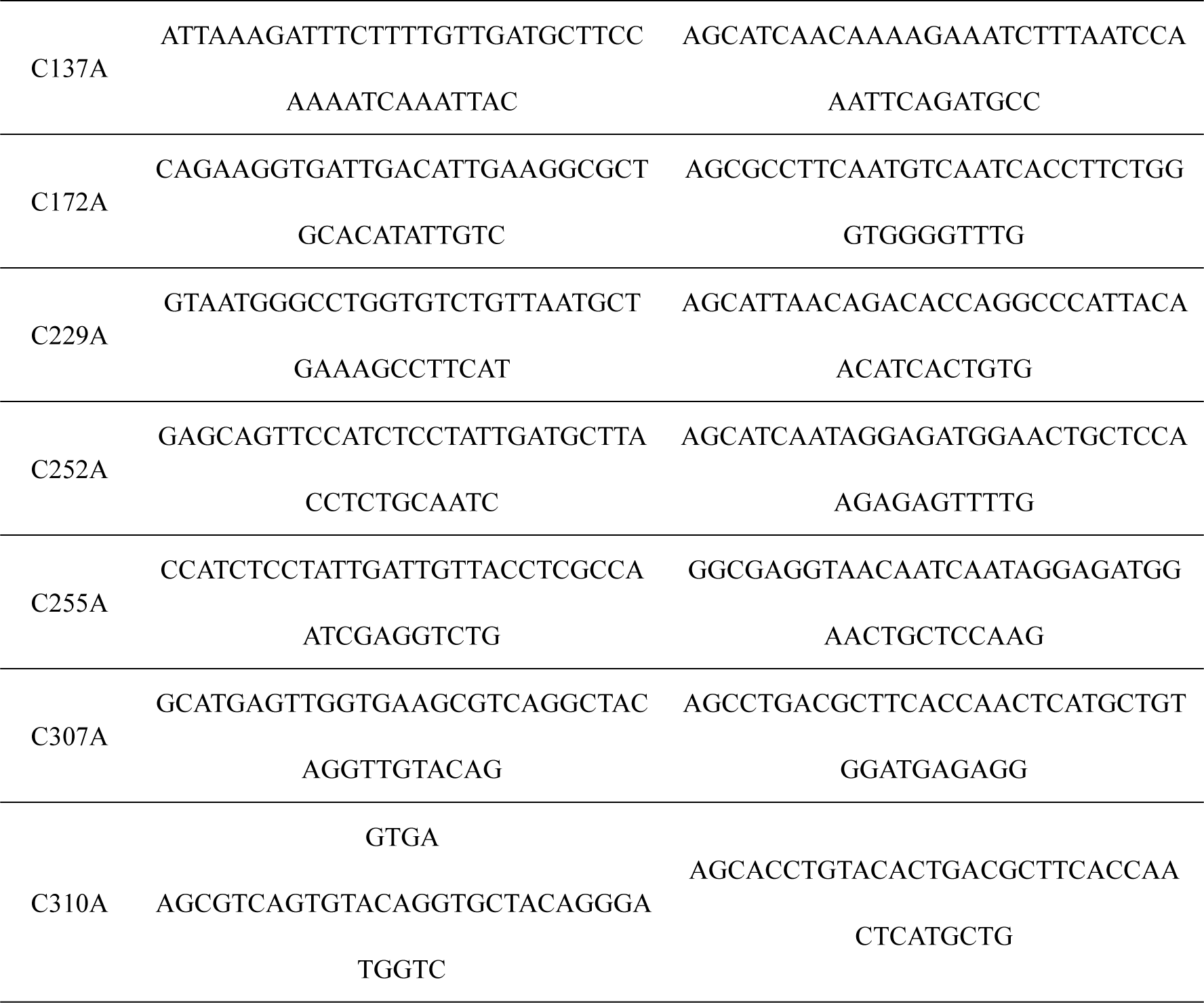
The construction of pIRES2-EGFP-hCTH-C70A、C84A、C109A、C137A、C172A、C229A、C252A、C255A、C307A and C310A.

